# The MIDAS domain of AAA mechanoenzyme Mdn1 forms catch bonds with two different substrates

**DOI:** 10.1101/2021.08.30.458255

**Authors:** Keith J. Mickolajczyk, Paul Dominic B. Olinares, Brian T. Chait, Shixin Liu, Tarun M. Kapoor

## Abstract

Catch bonds are a form of mechanoregulation wherein protein-ligand interactions are strengthened by the application of dissociative tension. Currently, the best-characterized examples of catch bonds are between single protein-ligand pairs. The essential AAA (ATPase associated with diverse cellular activities) mechanoenzyme Mdn1 drives two separate steps in ribosome biogenesis, using its MIDAS domain to extract the ubiquitin-like (UBL) domain-containing proteins Rsa4 and Ytm1 from ribosomal precursors. However, it must subsequently release these assembly factors to reinitiate the enzymatic cycle. The mechanism underlying MIDAS-UBL switching between strongly- and weakly-bound states is unknown. Here, we use single-molecule optical tweezers to investigate the force-dependence of MIDAS-UBL binding. Parallel experiments with Rsa4 and Ytm1 show that forces up to ~4 pN, matching the magnitude of force produced by AAA proteins similar to Mdn1, enhance the MIDAS domain binding lifetime up to tenfold, and higher forces accelerate dissociation. Together, our studies indicate that Mdn1’s MIDAS domain forms catch bonds with more than one UBL-substrate, and provide insights into how mechanoregulation may contribute to the Mdn1 enzymatic cycle during ribosome biogenesis.

## INTRODUCTION

Regulation of protein-ligand interactions is an essential organizational principle in cell biology, and is achieved through various means, including post-translational modifications (*e.g*. phosphorylation) and substrate exchange (*e.g*. GTP hydrolysis) (Alberts et al., 1994). More recently, mechanoregulation - or changes in protein-ligand interactions in response to applied mechanical forces - has garnered interest as another organizing principle. While most protein-protein interactions are known to become weaker when external forces are applied along the bond-axis, referred to as slip bonds, a few special cases have been identified where force increases bond lifetimes, referred to as catch bonds (Bell, 1978; Sokurenko et al., 2008). Catch bonds are hypothesized to be a widely used regulatory mechanism in the cell (Sokurenko et al., 2008), but force spectroscopy investigations to date have largely focused on cell adhesion and cytoskeleton proteins. Moreover, most known examples of catch bonds are for one protein interacting with one specific ligand.

Ribosome biogenesis is a complex, multi-step process involving hundreds of trans-acting protein and RNA factors, including ATP-consuming mechanoenzymes (Frazier et al., 2021; Kressler et al., 2010). One essential mechanoenzyme is Mdn1, a member of the AAA (ATPase associated with diverse cellular activities) superfamily. Mdn1 is a large (>500 kDa) multi-domain enzyme; at its N-terminus are six AAA domains, which fold into a pseudohexameric ring, followed by a linker including both a structured (~1,800 amino acids) and a likely non-structured (~517 amino acids) region, and finally a C-terminal (290 amino acid) MIDAS domain (**Figure 1A**) (Garbarino and Gibbons, 2002). Mdn1 binds and extracts the ubiquitin-like (UBL) domain-containing assembly factors Ytm1 and Rsa4, which are embedded in pre-60S (large) ribosomal precursors at different stages of maturation, via its MIDAS domain (Bassler et al., 2010; Ulbrich et al., 2009). Recent structural work revealed that the MIDAS domain docks onto the AAA ring in the context of pre-60S binding, and forms a tripartite connection in which it is bound to Rsa4 (Chen et al., 2018; Kater et al., 2020; Sosnowski et al., 2018). However, we do not understand how the MIDAS-UBL binding, which must be strong for assembly factor removal, switches to a more weakly-bound state, such that the UBL proteins can be subsequently released.

**Figure 1.**
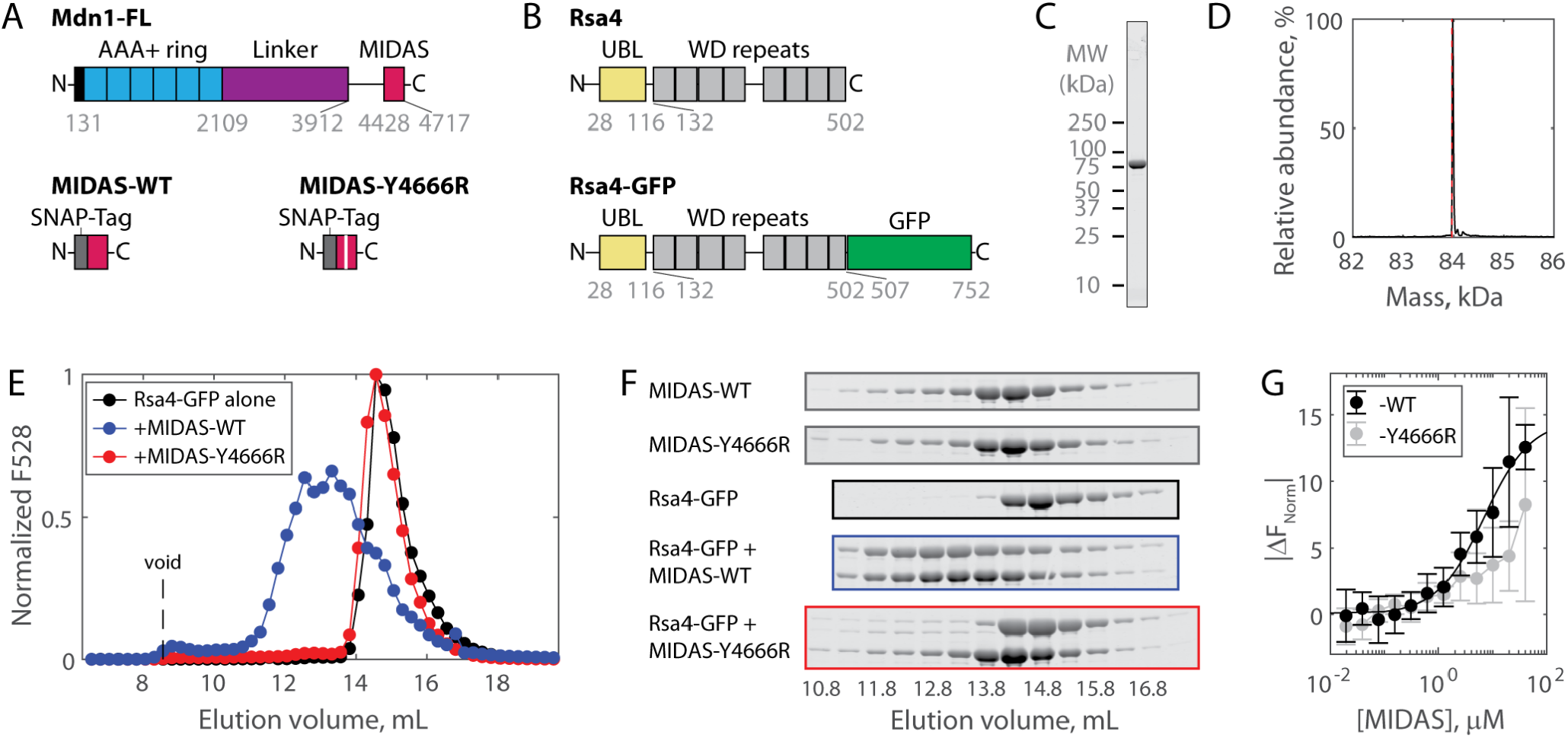
Solution measurements of MIDAS-Rsa4 interaction. **(A)** Domain diagrams (drawn to scale) of full-length Mdn1 (top), and the SNAP-tagged MIDAS constructs (bottom). **(B)** Domain diagrams (drawn to scale) highlighting the position of the UBL and WD repeat domains within the assembly factor Rsa4, as well as the added green fluorescent protein (GFP) tag. **(C)** SDS-PAGE gel (Coomassie staining) of final purified Rsa4-GFP. See also **Figure S1A**. **(D)** Native mass spectrometry analysis of Rsa4-GFP. Expected mass (with loss of N-terminal methionine) 83.962 kDa (red dotted line), measured mass 83.981±0.001 kDa (mean ± standard deviation, SD). Full spectrum in **Figure S1B**. **(E)** Elution profile (monitored by GFP fluorescence) of Rsa4-GFP (40 μM prior to injection) either alone or pre-mixed with MIDAS-WT or MIDAS-Y4666R (no GFP label; 60 μM) on a Superdex 200 Increase size exclusion column. **(F)** SDS-PAGE gels (Coomassie staining) corresponding to the elution profiles shown in panel E. See also **Figure S2**. **(G)** Microscale thermophoresis data showing the binding of MIDAS protein to Rsa4-GFP (50 nM). All data shown as mean ± SD for n=4 independent experiments including at least two separate preparations of each protein used. The MIDAS-WT data were fitted to a binding isotherm (black curve). MIDAS-Y4666R data connected by lines to guide the eye.

The Mdn1 MIDAS domain bears structural homology to the integrin α I domain (Garbarino and Gibbons, 2002). In solution, this integrin domain has a weak (~1 mM) affinity for its ligands (Shimaoka et al., 2001). Remarkably, this affinity can be enhanced by about an order of magnitude by applied force (Astrof et al., 2006; Kong et al., 2009; Shimaoka et al., 2003). Although a UBL domain is structurally distant from extracellular integrin ligands, it is possible that Mdn1 may similarly use mechanoregulation for its function. Consistent with this idea, Brownian dynamics simulations have shown that docking of the MIDAS domain onto the AAA ring stretches the unstructured portion of the linker such that 1-2 pN of tension is generated (Mickolajczyk et al., 2020). This tension would be transmitted along the MIDAS-UBL bond axis, and its magnitude may even be enhanced by ATP-dependent motions propagated from the AAA ring (Chen et al., 2018; Ulbrich et al., 2009). However, no studies to date have tested the hypothesis that the Mdn1 MIDAS domain can form catch bonds with its substrates.

In the current work, we investigate the mechanoregulation of the interaction between Mdn1’s MIDAS domain and two UBL domain-containing assembly factors. Using bulk assays, we find that MIDAS-Rsa4 and -Ytm1 affinity is weak (≥7 μM). Using a newly-developed optical tweezers “force jump” assay, we find that the Mdn1 MIDAS domain forms a catch bond with both Rsa4 and Ytm1, with bond lifetimes extending ~tenfold at ~4 pN tension. These measurements show that mechanical forces are sufficient for modulating Mdn1-substrate binding, and suggest that catch bonds play a key role in regulating Mdn1-driven steps in ribosome biogenesis.

## RESULTS

### Solution measurements of MIDAS-Rsa4 binding

To investigate the regulation of Mdn1 binding to its UBL domain-containing pre-ribosomal substrate Rsa4, we first sought to characterize binding *in vitro*. The Mdn1 MIDAS domain (amino acids 4381-4717) with an N-terminal SNAP tag was expressed in *E. coli* as previously (**Figure 1A**) (Mickolajczyk et al., 2020). Both wild type MIDAS domain (MIDAS-WT) and MIDAS domain harboring a point mutation known to disrupt UBL domain binding (MIDAS-Y4666R) were generated (Ahmed et al., 2019). Full-length Rsa4 with a C-terminal GFP (Rsa4-GFP; **Figure 1B**) was expressed in insect cells and purified using affinity, ion exchange, and size exclusion chromatography (see **methods**). Purity was assessed by SDS-PAGE (**Figure 1C, S1A**). Rsa4-GFP was confirmed to be a monomer of the correct molecular weight both by native mass spectrometry (to within 20 Da; **Figure 1D, S1B**) and by mass photometry (to within 2 kDa; **Figure S1C**).

Binding of the Mdn1 MIDAS domain to Rsa4-GFP was first assessed by size exclusion chromatography coelution assays. Rsa4-GFP (40 μM prior to injection) by itself eluted as a single peak at ~14.5 mL, as measured by GFP fluorescence (**Figure 1E**) and SDS-PAGE (**Figure 1F**, black; uncropped gels in **Figure S2**). MIDAS-WT alone and MIDAS-Y4666R alone (60 μM prior to injection) did not produce a fluorescence signal, but were each seen to elute at ~14.3 mL by SDS-PAGE (**Figure 1F**, gray). Mixing Rsa4 with MIDAS-WT before injection led to a shift in the fluorescence profile towards a higher molecular weight, with an overlapping elution profile consistent with binding (**Figure 1E-F**, blue). Mixing Rsa4 with MIDAS-Y6664R before injection did not lead to a shift in the fluorescence profile or to apparent coelution by SDS-PAGE (**Figure 1 E-F**, red), suggesting that the coelution with MIDAS-WT is due to specific binding.

We next sought to determine the affinity for the Mdn1 MIDAS domain for Rsa4-GFP using microscale thermophoresis (MST). Titrating MIDAS-WT against Rsa4-GFP (50 nM) produced changes in the normalized fluorescence signal upon heating (|ΔF_Norm_|) that could be fitted to a binding isotherm with K_D_= 6.9 ± 2.0 μM (fit ± 95% confidence intervals, CI) (**Figure 1G**). Titrating MIDAS-Y4666R led to smaller |ΔF_Norm_| values that could not be fitted, consistent with weak binding. Together, these results show that while the Mdn1 MIDAS domain can bind to Rsa4 in solution, the binding affinity is weak (~7 μM).

### Single-molecule measurements of MIDAS-Rsa4 binding under applied load

To investigate the force-dependence of the MIDAS-UBL domain interaction, we developed a single-molecule optical tweezers “force jump” assay (**Figure 2A**). In this assay, two double-stranded (dsDNA) handles are attached to two polystyrene beads - one held in a micropipette by suction and the other held by an optical trap - and connected using a single-stranded (ssDNA) bridge. SNAP-tagged MIDAS and GFP nanobody proteins are covalently bound to ssDNA oligonucleotides (**Figure 2B**), which are then annealed to the bridge strand (**Figure 2C**). Rsa4-GFP in solution can bind to the GFP nanobody on the bridge construct. With the tethered assembly in place, the optical trap, in constant force mode, can be rapidly switched (*i.e*. force jumped) between a low and a high constant force. At the high force, MIDAS and Rsa4-GFP are physically separated and cannot bind (**Figure A** state 1). At the low force (**Figure 2A** state 2) the bridge strand is relaxed and MIDAS and Rsa4 come into close enough proximity to bind. Should the proteins bind while at the low force, an intermediate position can be read out when the system is jumped to high force (**Figure 2A** state 3). This intermediate position informs on the lifetime of the MIDAS-Rsa4 interaction. The distance traveled between the intermediate and final high-force position (Δx) should depend on the length of the single-stranded region of annealed bridge construct. We thus performed our assays with two bridge constructs of different lengths (55 and 70 nt; **Figure 2C**). For both bridge constructs, we ensured that only a single “tether” was drawn between each pair of beads by examining the shape of the force-extension curve (**Figure 2D**).

**Figure 2.**
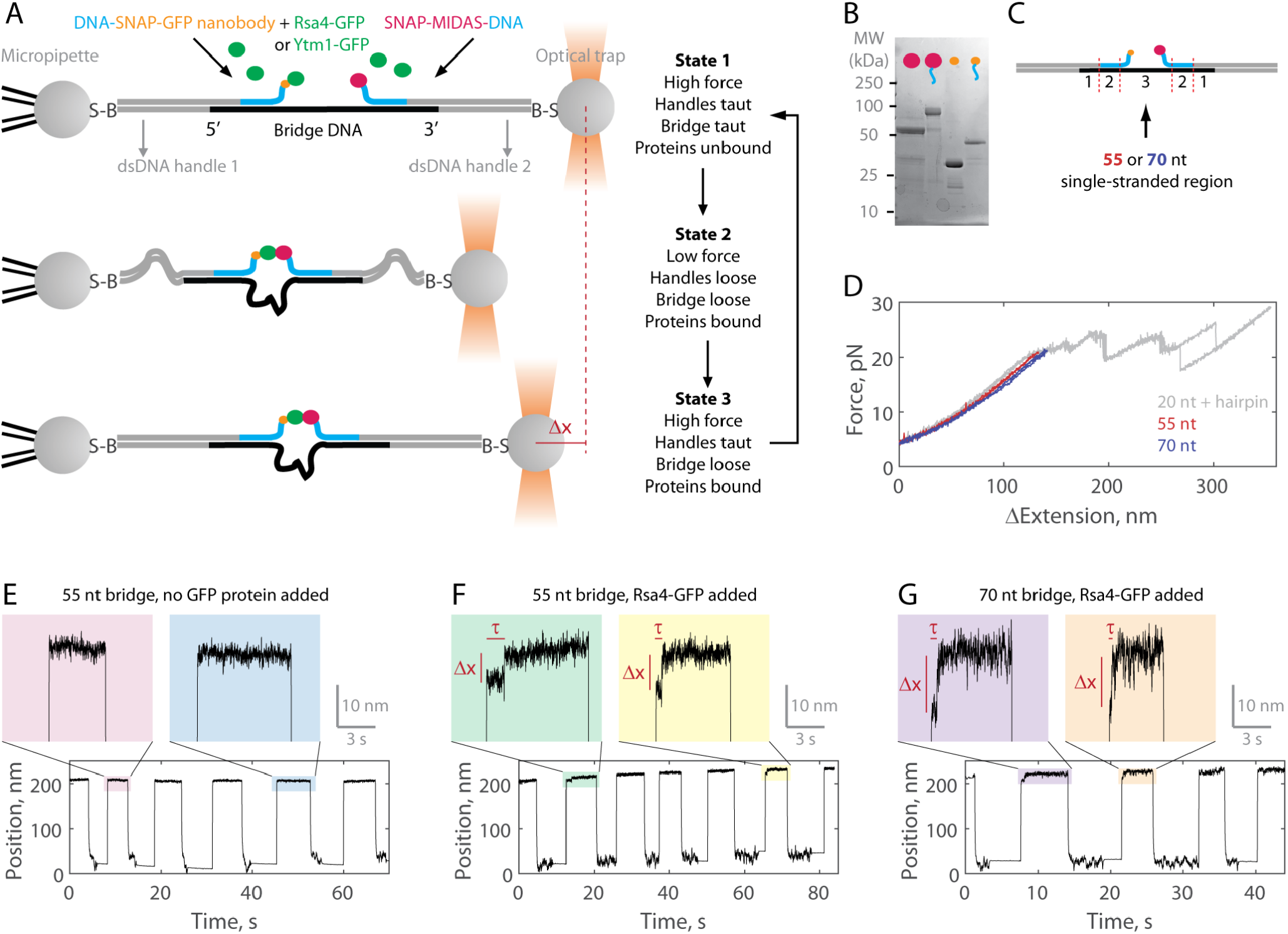
Single-molecule assay for measuring the MIDAS-Rsa4 interaction under load. **(A)** Single-molecule optical tweezers “force jump” assay design. Two 1.5 kilobase-pair double-stranded (dsDNA) handles are attached to 2.1 μm beads via biotin-streptavidin (B-S) linkage. The handles have 31-33 nucleotide (nt) overhangs, to which a single-stranded (ssDNA) “bridge” is annealed. A copy of GFP nanobody and MIDAS protein, each conjugated to a DNA oligonucleotide (blue) are then annealed to the bridge. Rsa4-GFP, free in solution, can bind to the GFP nanobody. Dissociation of the MIDAS-UBL interaction under load can be read out as a positional change (Δx) when the optical trap (in constant force mode) is switched from a low to a high applied force. **(B)** SDS-PAGE gel (Coomassie staining) showing the covalent attachment of ssDNA oligonucleotides to the SNAP-tagged MIDAS-WT (pink) and SNAP-tagged GFP nanobody (orange). **(C)** Sections of the DNA bridge. Section 1 (31-33 nt) anneals to the dsDNA handles, section 2 (30 nt) anneals the protein-bound DNA oligonucleotides, and section 3 remains single-stranded. Different lengths of section 3 (55 or 70 nt) are expected to produce Δx events of different magnitude. **(D)** Example force-extension curves of the DNA handles connected by the 55 nt (red) and 70 nt (blue) bridge. Also shown is a hairpin (gray) which anneals to the dsDNA handles with a 20 nt region leftover. The distinct unfolding/refolding pattern at loads above 20 pN enables identification of a single “tether” between the two beads. **(E)** Example force jump data (0.5 and 6 pN low and high force) on the 55 nt bridge with all components added except a GFP-labeled assembly factor. In some instances, force-feedback was released at the low force level to reduce large fluctuations. Only two position levels were observed. **(F)** Example force jump data on the 55 nt bridge with Rsa4-GFP (20 nM) added. On some jumps (highlighted), an intermediate position is observed. **(G)** Similar to panel F, with the 70 nt bridge construct.

Example data generated in the force jump assay are shown in **Figure 2 E-G**. Here we held a constant force of 0.5 pN for at least 5 seconds before jumping to a constant force of 6 pN, also held for at least 5 seconds. When no GFP-tagged assembly factor was present, only two positional states were detected (here using the 55 nt bridge construct; **Figure 2E**). When Rsa4-GFP was added (20 nM) an intermediate position appeared on some of the jumps, consistent with MIDAS-Rsa4 binding (**Figure 2F**). For these intermediate positions, we quantified both a bond lifetime (τ) and a distance change (Δx) using a two-state hidden Markov model algorithm (see **Methods**). We next made measurements using the 70 nt bridge construct, and again saw intermediate positions, but with larger Δx magnitudes (**Figure 2G**). For both the 55 and 70 nt bridge constructs, intermediate positions were not seen in the negative control (**Figure S3A**). Altogether, these data exemplify single-molecule measurements of MIDAS-Rsa4 binding under applied forces.

### Force-dependence of MIDAS-Rsa4 binding

We next sought to measure the force-dependence of binding by quantifying bond lifetimes (τ) of MIDAS and Rsa4-GFP on both bridge constructs (55 and 70 nt) at multiple high-force levels (hereafter referred to as total applied force levels, F_Tot_). We pooled data for multiple (n=26-27) tethers that displayed intermediate positions (**Figure 2D**, **S3A**). We first measured MIDAS-WT with Rsa4 on the 55 nt bridge construct at 6 pN (**Figure 3A**). The distribution (n=33) could be fitted to a single exponential (appears linear on a semi-log plot), consistent with the kinetics of exit from a single bound state (Guo and Guilford, 2006). The inverse of the average bond lifetime is equivalent to the off-rate. As an additional control, we measured the bond lifetime of Rsa4-GFP with oligonucleotide-bound MIDAS-Y4666R, the weak-binding mutant (also run on the 55 nt tether construct with an F_Tot_ value of 6 pN). This data (n=29) could also be described with a single exponential. The substantially shorter bond lifetime distribution of MIDAS-Y4666R with Rsa4-GFP versus MIDAS-WT provides evidence that the binding events measured are specific.

**Figure 3.**
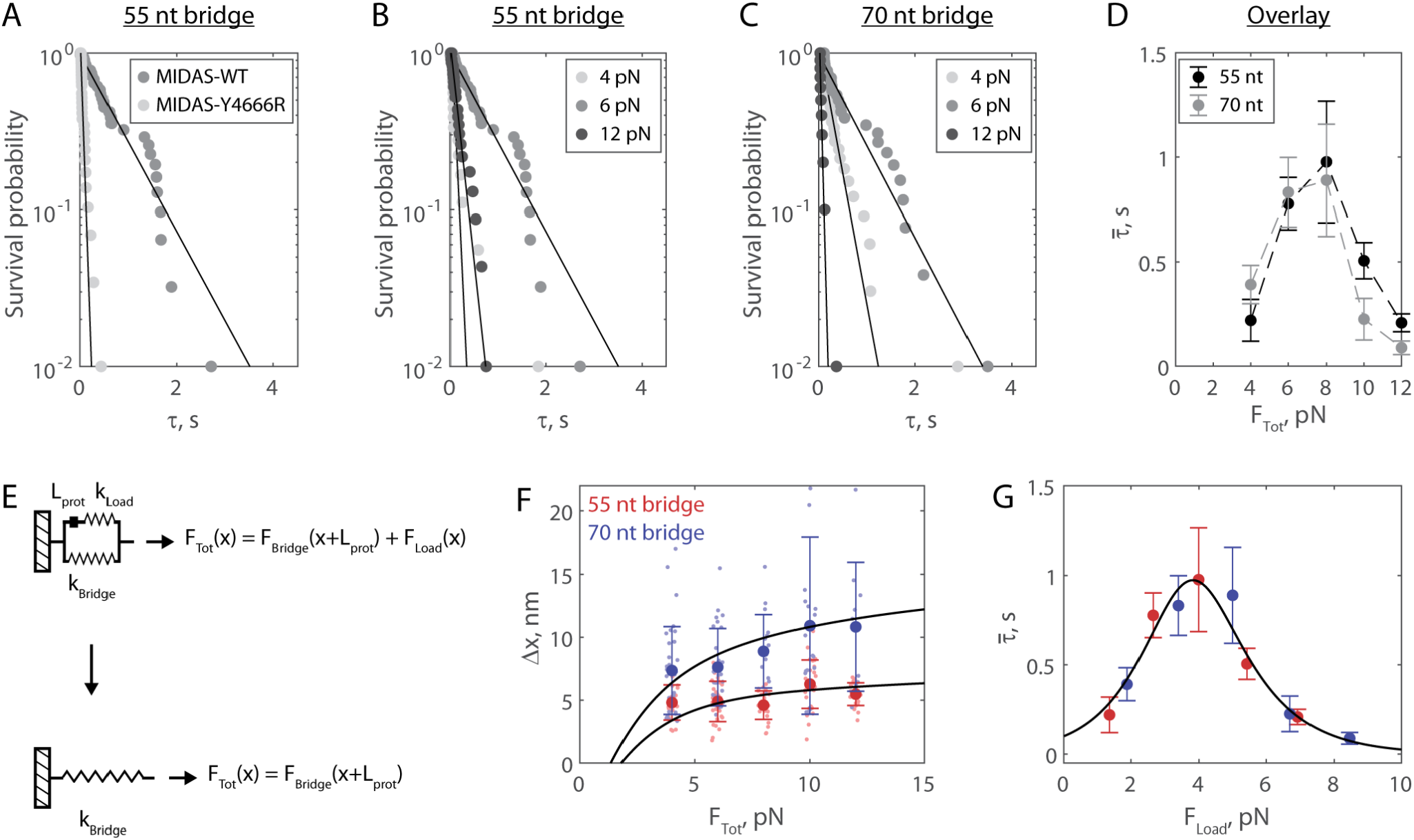
The Mdn1 MIDAS domain forms a catch bond with Rsa4. **(A)** Distributions of MIDAS-Rsa4 bond lifetimes (6 pN total force, 55 nt bridge construct) with either MIDAS-WT (n=31) or MIDAS-Y4666R (n=29). For all distributions, the final data point was moved from y=0 to y=0.01 to enable semilog plotting. Black lines show fits to a single exponential. **(B)** Distributions of MIDAS-Rsa4 bond lifetimes on the 55 nt bridge construct at 4 pN (n=18), 6 pN (n=31), and 12 pN (n=23) total applied force. **(C)** Distributions of MIDAS-Rsa4 bond lifetimes on the 70 nt bridge construct at 4 pN (n=33), 6 pN (n=26), and 12 pN (n=10) total applied force. **(D)** The average bond lifetime of MIDAS-Rsa4 binding as a function of total applied force. Data shown as mean ± standard error of the mean (SEM; n=10-33 measurements), with dotted lines to guide the eye. **(E)** Mechanical circuit model describing the force jump assay. When the proteins (of inextensible length L_prot_) are bound, force is partitioned between the top “loading” strand (two 12 nt single-stranded regions) and the bottom bridge strand. In the equations shown inset, x designates extension along the loading stand spring. **(F)** The magnitude of Δx for Rsa4-MIDAS interactions as a function of total applied force. Individual measurements shown as small data points, mean ± SD (n=10-33) shown in bold. Data generated with the 55 and 70 nt bridge constructs shown in red and blue, respectively. Black lines show output of the mechanical circuit model (see also **Figure S4**). **(G)** The average MIDAS-Rsa4 bond lifetime as a function of force applied across the proteins. Data generated with the 55 nt bridge construct shown in red and data generated with the 70 nt bridge construct shown in blue. Data points shown as mean ± SEM (n=10-33 measurements). Black curve shows fit (weighted by inverse SEM) to the catch-slip Bell model.

We next built distributions for the bond lifetime of Mdn1 MIDAS-WT with Rsa4-GFP at multiple values of F_Tot_ on both the 55 (**Figure 3B**) and 70 nt (**Figure 3C**) tether constructs (All distributions in **Figure S3B**, n=8-33). In each of these experiments, we observed that the distribution shifted from short to long bond lifetimes between 4 and 6 pN, and then back to short bond lifetimes at 12 pN. All measured distributions could be fitted to a single exponential (**Figure S3B**), indicating that applied force influenced the kinetics of dissociation, not the number of states from which dissociation could occur (Huang et al., 2017). Plotting the average bond lifetime 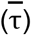 as a function of F_Tot_, we noted a chevron-shaped curve for both bridge constructs with minor differences at low and high F_Tot_ values (**Figure 3D, S3B**). The average bond lifetime increased between F_Tot_ values of 4-8 pN, and then decreased at F_Tot_ values of 10-12 pN. This stabilization of binding by external load up to a certain threshold (catch bond) contrasts the more common scenario where external force monotonically accelerates dissociation (slip bond).

### Mdn1 MIDAS and Rsa4 form a catch bond

In our experimental geometry, the total force is partitioned between the protein pair and the bridge DNA (**Figure 2A**). To derive the force applied to the protein bond from F_Tot_, we built a mechanical circuit model of the force jump assay (**Figure 3E, S4**). When MIDAS and Rsa4-GFP are bound, we model the system as two nonlinear springs in parallel. The top (loading) spring consists of the 12 nt sections of the oligonucleotides bound to SNAP-GFP nanobody and SNAP-MIDAS that do not anneal to the bridge DNA, as well as the relatively rigid folded proteins, which we model as inextensible (constant length L_prot_) at 8 nm (Ahmed et al., 2019). The bottom spring is the non-complementary ssDNA region of the bridge DNA (55 or 70 nt). Since the proteins are on the top spring, the bottom spring begins pre-stretched relative to the top spring by L_prot_. In this configuration, the total force applied by the optical trap (F_Tot_) is partitioned over the two springs (F_Load_ and F_Bridge_). When MIDAS and Rsa4-GFP dissociate from one another, the entirety of F_Tot_ is put onto the bridge DNA, leading to further stretching (Δx). We modeled the nonlinear springs in the system using the worm-like chain equation, a model for the elasticity of DNA (see **Methods**). An overlay of experimental measurements of Δx with output of the mechanical circuit model is shown in **Figure 3F**. We note that the model output (black lines) is not fitted to or constrained by the experimental data. The excellent agreement between theory and experiment provides further validation of the force jump assay and the mechanical circuit model.

We next used the mechanical circuit model to combine the bond lifetime data generated using the 55 and 70 nt bridge constructs (**Figure 3D**) and plot them as a function of F_Load_, the force actually placed on the protein-protein interaction (**Figure 3G**). The combined data smoothed the chevron shape, and could be fitted to the “catch-slip” application of the Bell model (Barsegov and Thirumalai, 2005; Evans et al., 2004; Evans and Ritchie, 1997; Guo and Guilford, 2006):

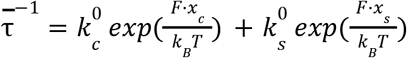

Where 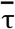 is the average bond lifetime and subscripts c and s refer to the catch and slip pathways, respectively. Here the catch pathway dominates at low forces and the slip pathway dominates at high forces, respectively generating the rise and fall of the chevron shape. We estimated parameters 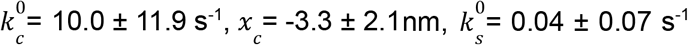, and *x_s_* = 2.8 ± 1.3 nm (fit ± 95% CI) for MIDAS with Rsa4-GFP. The similar values of the distance parameters x_c_ and x_s_ in both cases give rise to a roughly symmetric rise and fall in force-dependent bond lifetime, similar to the case of the P-selectin complex with sPSGL-1 (Barsegov and Thirumalai, 2005). Additionally, this fit allowed us to determine the critical force at which bond lifetime is maximized, which we estimated to be 3.8 pN. Overall, this analysis shows that the Mdn1 MIDAS domain forms a catch bond with Rsa4.

### Mdn1 MIDAS also forms a catch bond with Ytm1

We next tested whether Mdn1 MIDAS catch bond behavior is specific for Rsa4, or more general with another UBL domain-containing ribosome assembly factor. Mdn1 has been proposed to also remove Ytm1, from the pre-60S particle (Ahmed et al., 2019; Bassler et al., 2010). However, the *S. pombe* Rsa4 and Ytm1 UBL domains only have 43.0% sequence similarity. Ytm1-GFP (**Fig. 4A**) was generated in insect cells similarly to Rsa4-GFP. Ytm1-GFP ran with an apparent double-banding pattern (**Figure S1A**), however the protein was assessed to be homogenous, monomeric, and centered around the expected molecular weight both by native mass spectrometry (to within 20 Da; **Figure S1B**) and mass photometry (to within 1 kDa; **Figure S1C**). We ran size exclusion chromatography assays, and as with Rsa4-GFP, coelution was seen for Ytm1-GFP with MIDAS-WT but not with MIDAS-Y4666R (**Figure 4B-C, S2**). We also used MST to assess MIDAS-Ytm1-GFP binding. Titrating MIDAS-WT against Ytm1-GFP (50 nM) led to larger |ΔF_Norm_| values than titrating MIDAS-Y4666R, but the data could not be fitted to a binding isotherm in either case (**Figure 4D**). Hence. MIDAS and Ytm1-GFP can bind *in vitro*, but the binding in solution is even weaker than that of MIDAS and Rsa4-GFP.

**Figure 4.**
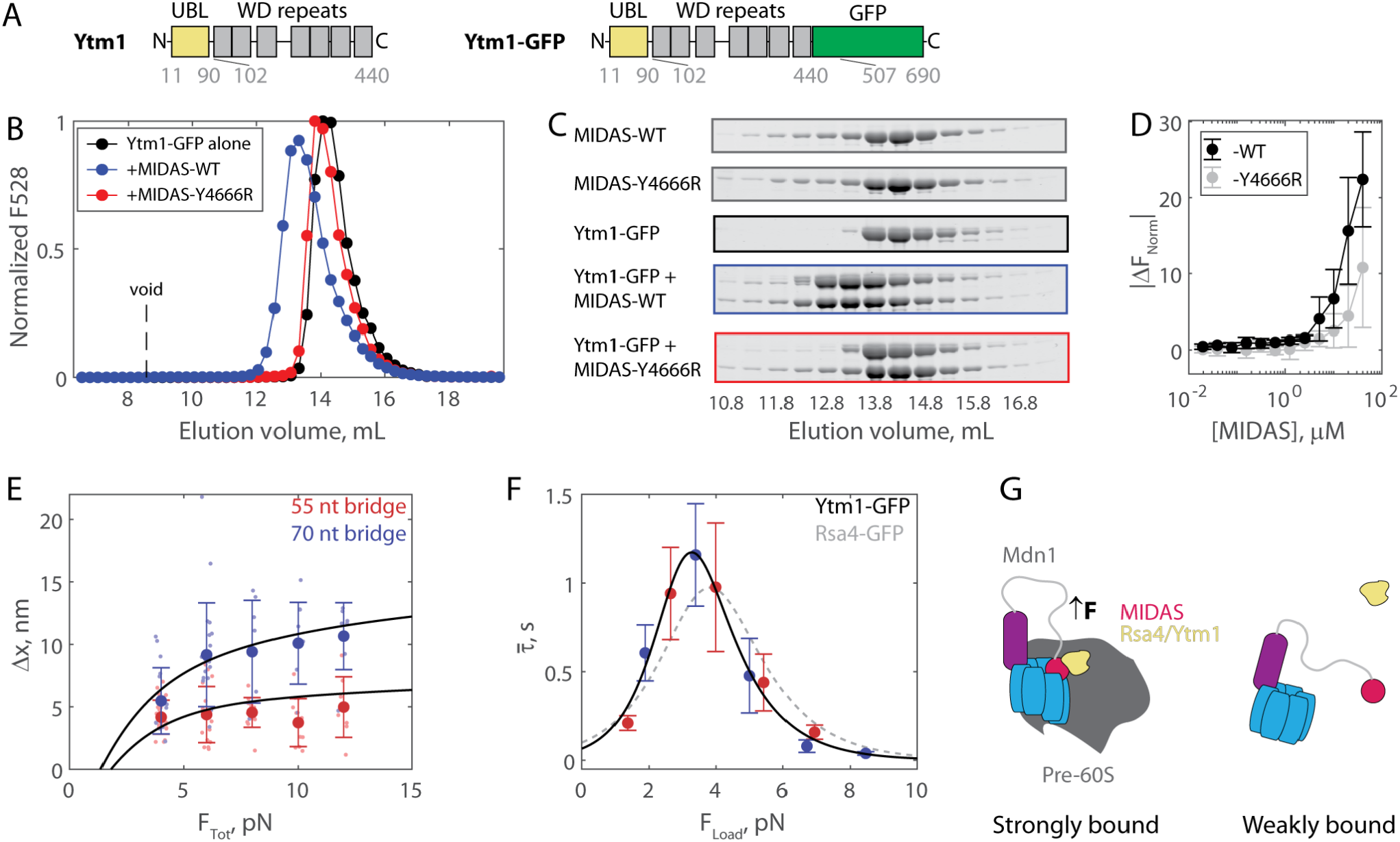
The Mdn1 MIDAS domain also forms a catch bond with Ytm1. **(A)** Domain diagrams (drawn to scale) highlighting the position of the UBL and WD repeat domains within the assembly factor Ytm1, as well as the added green fluorescent protein (GFP) tag. **(B)** Elution profile (monitored by GFP fluorescence) of Ytm1-GFP (40 μM prior to injection) either alone or pre-mixed with MIDAS-WT or MIDAS-Y4666R (no GFP label; 60 μM) on a Superdex 200 Increase size exclusion column. **(C)** SDS-PAGE gels (Coomassie staining) corresponding to the elution profiles shown in panel B. **(D)** Microscale thermophoresis data showing the binding of unlabeled MIDAS protein to Ytm1-GFP (50 nM). All data shown as mean ± SD (n=4). Data points connected by lines to guide the eye. **(E)** The magnitude of Δx for Ytm1-MIDAS interactions as a function of total applied force. Individual measurements shown as small data points, mean ± SD (n=8-22) shown in bold. Black lines show output of the mechanical circuit model. Example traces with Ytm1-GFP in the force jump assay shown in **Figure. S3C**. **(F)** The average bond lifetime of MIDAS-Ytm1 binding as a function of force applied across the proteins. Data generated with the 55 nt bridge construct shown in red and data generated with the 70 nt bridge construct shown in blue. Data points shown as mean ± SEM (n=8-22 measurements). All distributions in **Figure S3D**. Black curve shows fit (weighted by inverse SEM) to the catch-slip Bell model. Gray dotted line shows the fit for MIDAS-Rsa4 binding (**Figure 3G**) for comparison. **(G)** MIDAS-UBL catch bond in ribosome biogenesis. Force generated in the AAA ring of Mdn1 (blue) must be transmitted through the MIDAS domain to mechanochemically remove an assembly factor, Rsa4 or Ytm1 (yellow), from the pre-60S particle. The MIDAS domain binds Rsa4 or Ytm1 strongly when Mdn1 is applying a force, and weakly after the proteins dissociate from the pre-60S particle.

We next ran the force jump assay with Ytm1-GFP present instead of Rsa4-GFP. We measured bond lifetimes and Δx values using both the 55 and 70 nt bridge constructs, and measured a force-dependence of binding (**Figure S3C-D**). Measurements of Δx made with Ytm1-GFP present fell along the expected curves based on the mechanical circuit model (without any free parameter), similar to Rsa4-GFP (**Figure 4E**). Plotting the mean bond lifetime as a function of F_Load_, the force experienced by the proteins in the force jump assay, we found that Mdn1 MIDAS again forms a catch-slip type bond with Ytm1-GFP (**Figure 4F**). The fitted parameters were similar to those of Rsa4-GFP: 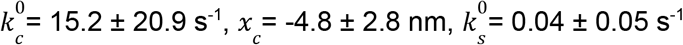, and *x_s_* = 3.2 ± 0.9 nm (fit ± 95% CI). The critical force where binding lifetime is maximized was 3.3 pN, versus 3.8 pN for Rsa4-GFP. Overall, these results show that the Mdn1 MIDAS domain forms a catch bond with two different UBL domain-containing proteins.

## DISCUSSION

In the current work, we investigate the regulation of binding between the Mdn1 MIDAS domain and the UBL domain-containing ribosome assembly factors Rsa4 and Ytm1. We show that the Mdn1 MIDAS domain has weak affinity for Rsa4 and Ytm1 in solution (≥7 μM). Using a single-molecule optical tweezers force jump assay developed for this study, we show that Mdn1 MIDAS forms a catch bond with both Rsa4 and Ytm1, with bond lifetimes increasing by an order of magnitude between 0-4 pN of applied tension, and decreasing back to baseline between 4-10 pN. Together, our findings provide insights into how forces may directly regulate Mdn1-driven steps in the nucleolus and nucleoplasm during ribosome biogenesis.

Based on our findings, we propose a model where the Mdn1 MIDAS-UBL catch bond plays a key role in assembly factor processing (**Figure 4G**). Mdn1 approaches the pre-60S particle and binds via its AAA ring, at which point the MIDAS domain both docks onto the AAA ring and binds the UBL-domain substrate (Chen et al., 2018; Kater et al., 2020; Sosnowski et al., 2018). In this MIDAS-docked state, Mdn1’s unstructured linker is stretched, building 1-2 pN of tension that is propagated along the MIDAS-UBL bond axis (Mickolajczyk et al., 2020). While the directionality of Mdn1 force generation has not been determined, one possibility is that ATPase activity in the AAA ring adds more tension to the unstructured linker, which would be propagated to the MIDAS-UBL interaction (**Figure 4G**), and thereby activate the MIDAS-UBL catch bond. The MIDAS domain remains bound to the UBL domain until the assembly factor dissociates from the pre-60S particle. Subsequently, the MIDAS domain undocks from the AAA ring, relieving tension across the bond with the UBL domain, deactivating the catch bond and facilitating assembly factor release. Ytm1 and Rsa4 removal occur at separate stages of pre-60S maturation, in the nucleolus and nucleus, respectively (Bassler et al., 2010; Kressler et al., 2012). We propose that our model holds for both these scenarios. We also note that free copies of Ytm1 and Rsa4 are present in the nucleolus and nucleus (Kressler et al., 2012); the Mdn1 catch bond thus also prevents Mdn1-Rsa4/Ytm1 binding while off the pre-60S, which would compete away usable copies of each protein.

It is interesting to note that Mdn1 processes UBL domain-containing proteins. The UBL domains of Rsa4 and Ytm1 have low sequence similarity (43.0% in *S. pombe*), but a high level of structural similarity (Ahmed et al., 2019; Romes et al., 2016). There are numerous (>10 in *S. pombe*, >60 in humans) other UBL domain-containing proteins with similar folds to Ytm1/Rsa4-UBL (Collins and Goldberg, 2020; Hartmann-Petersen and Gordon, 2004). Moreover, UBL domains can be covalently conjugated to other proteins by a cascade of UBL-specific enzymes (Schulman, 2011; Streich and Lima, 2014). In addition, Mdn1 in mammalian cells has been shown to be targeted to SUMOylated substrates (Raman et al., 2016), where SUMO and UBL domains also share structural similarity. While no Mdn1 MIDAS binding partners other than Rsa4 and Ytm1 have been confirmed to date, it is noteworthy that Mdn1 was recently implicated to also have a role in heterochromatic RNA clearance (Shipkovenska et al., 2020). Therefore, we speculate that Mdn1 may also process UBL- or SUMO-domain containing proteins other than Rsa4 and Ytm1 in the cell. Based on the close similarity of the Rsa4 and Ytm1 catch bonds measured, we predict that the Mdn1 MIDAS domain would also form catch bonds with these putative substrates.

The Mdn1 MIDAS domain bears structural homology to the integrin MIDAS domain (Chen et al., 2018; Garbarino and Gibbons, 2002). Integrin has also been shown to form catch bonds, albeit with a fragment of its substrate (containing the RGD consensus sequence) rather than with two different full-length, folded proteins (Astrof et al., 2006; Kong et al., 2009; Puklin-Faucher et al., 2006; Shimaoka et al., 2003). Interestingly, the critical force at which bond lifetime is maximized is ~10-30 pN for integrin (Chen et al., 2011; Kong et al., 2009), a force-range relevant for meso-scale cell adhesions, in contrast to ~3-4 pN for Mdn1 MIDAS (**Figure 3G, 4F**), matching the forces that dynein, the AAA protein with highest sequence similarity to Mdn1, is capable of producing (4.3 pN) (Belyy et al., 2016). We therefore suggest that MIDAS domains can be tuned to form catch bonds at forces ranging from the single motor to the multi-motor regime, as needed for specific cellular contexts. Towards this end, numerous AAA motor proteins including CbbQ (Sutter et al., 2015; Tsai et al., 2019, 2015), Vwa8 (Luo et al., 2017), magnesium chelatase (Fodje et al., 2001), and others (Iyer et al., 2004), as well as other proteins with known mechanical activation such as von Willebrand Factor (Fredrickson et al., 1998; Kim et al., 2010; Ruggeri and Ware, 1993), either contain or couple their activities with MIDAS domains. Further studies are needed to investigate the mechanoregulation of these MIDAS domains binding to their substrates.

Catch bonds are an example of how mechanical forces can directly regulate essential protein-ligand interactions. As opposed to other forms of regulation, such as phosphorylation, mechanoregulation is nearly instantaneous, and reversible without the need for accessory enzymes. Examples of known catch bonds range from cell adhesion through FimH (Thomas et al., 2002), selectins (Marshall et al., 2003) and vinculin (Huang et al., 2017), to outside-in signaling through integrins (Kong et al., 2009) and notch-jagged (Luca et al., 2017), to motor proteins including dynein (Kunwar et al., 2011) and myosin (Guo and Guilford, 2006), and even to kinetochore attachments during mitosis (Akiyoshi et al., 2010). Our data uncovers catch bonds between proteins involved in ribosome biogenesis, and, in contrast to these known examples, between a protein and more than one substrate. We anticipate that further examples of proteins that form catch bonds with more than one substrate will be identified in the future, and the force jump assay developed here can serve as a valuable tool in this effort.

## METHODS

### Protein expression and purification

The SNAP-MIDAS construct, as reported previously (Mickolajczyk et al., 2020), was generated by subcloning Mdn1 aa4381-4717 into pSNAP-tag(T7)-2 (NEB N9181S) downstream of the SNAP tag. A N-terminal 6xHis tag and a Tobacco Etch Virus (TEV) protease site were added upstream of the SNAP tag. The Y4666R construct was generated using basic restriction cloning, verified by Sanger Sequencing. Both MIDAS constructs were expressed and purified using the same protocol. MIDAS proteins were expressed in *E. coli* Rosetta (DE3) pLysS cells (Merck 70954) grown in Miller’s LB medium (Formedium LMM105). Expression was induced at A600 = 0.6–0.8 with 0.5 mM IPTG (Goldbio), and the cultures were grown at 18°C for 16 hours. Cultures were pelleted and resuspended in lysis buffer (50 mM HEPES [pH 7.5], 150 mM NaCl, 1 mM MgCl_2_, 10% w/v glycerol, 20 mM imidazole, 5 mM 2-mercaptoethanol, 1 mM PMSF, 3 U/mL benzonase, 1X Roche complete protease inhibitor without EDTA). All purification steps were carried out at 4° C. Cells were lysed using an Emulsiflex-C5 homogenizer (Avestin, 4 cycles at ~10,000 psi), and the crude lysate was centrifuged at 55,000 rpm in a Type 70 Ti rotor (Beckman Coulter) for 30 minutes. The supernatant was filtered through a 0.22 μm Millex-GP PES membrane (Millipore SLGP033RS) and loaded onto a HisTrap HP column (Cytiva 17524701) pre-equilibrated with wash buffer (50 mM HEPES [pH 7.5], 150 mM NaCl, 1 mM MgCl_2_, 10% w/v glycerol, 20 mM imidazole, 5 mM 2-mercaptoethanol). The column was washed with 25 mL of wash buffer and eluted with wash buffer plus 400 mM imidazole. The eluent was treated with TEV protease (~0.1 mg/ml) and dialyzed against 1 L of dialysis buffer (20 mM HEPES [pH 7.5], 100 mM NaCl, 1 mM MgCl_2_, 10% w/v glycerol, 5 mM 2-mercaptoethanol) for 3 hours at 4°C. The supernatant was then loaded onto a HiTrap Q HP column (Cytiva 29051325) and eluted over a gradient of low salt (same as dialysis buffer) and high salt (dialysis buffer with 1 M NaCl) buffers. Relevant factions were located by SDS-PAGE, pooled, and concentrated using a 30 kDa cutoff Amicon Ultra-4 centrifugal filter (Millipore UFC803008). The samples were then loaded onto a Superdex 200 Increase 10/300 GL column (Cytiva 28990944) in binding assay buffer (50 mM Tris [pH 7.5], 80 mM NaCl, 5 mM MgCl_2_, 5% w/v glycerol, 0.002% Tween-20, 1 mM DTT). Concentration was determined using the colorimetric Bradford assay (Bio-Rad 5000006).

The plasmid containing the SNAP-tagged GFP nanobody was purchased on Addgene (#82711). This plasmid was transformed into Rosetta cells and prepared as described above. The same protocols and buffers were used up through elution from the HisTrap column. The His-eluent was concentrated using a 30 kDa cutoff Amicon Ultra-4 centrifugal filter (Millipore UFC803008), filtered using a 0.22 μm Millex-GP PES membrane (Millipore SLGP033RS), and loaded onto the Superdex 200 Increase 10/300 GL column (Cytiva 28990944) in storage buffer (20 mM HEPES [pH 7.5], 150 mM NaCl, 1 mM MgCl_2_, 3% w/v glycerol, 1 mM DTT).

Rsa4 and Ytm1 were amplified from a *S. pombe* mitotic cDNA library (Cosmo Bio 02-703) and cloned into pAceBa1c (Geneva Biotech) with C-terminal GFP, TEV, and 6xHis tags Recombinant baculoviruses were generated using the Bac-to-Bac system (Thermo Fisher). High Five cells (Thermo Fisher B85502) were grown to ~3.0 million cells/ml in Express Five SFM (Thermo Fisher 10486025) supplemented with antibiotic-antimycotic (Life Technologies 15240-062) and 16 mM L-glutamine (Life Technologies 25030-081) prior to infection with P3 viral stocks at a 1:50 virus:media ratio. Cells were cultured in suspension (27°C, shaking at 115 RPM) for 48 hours prior to harvesting.

Rsa4-GFP purification was carried out at 4° C. Cells were lysed using a Dounce homogenizer (Thomas Scientific) in ~25 mL of lysis buffer (50 mM HEPES [pH 6.5], 400 mM NaCl, 1 mM MgCl_2_, 10 % w/v glycerol, 20 mM imidazole, 5 mM 2-mercaptoethanol, 0.2 mM ATP, 1 mM PMSF, 3 U/mL benzonase, 1X Roche complete protease inhibitor without EDTA) per L of initial cell culture. The crude lysate was centrifuged at 55,000 rpm in a Type 70 Ti rotor (Beckman Coulter) for 1 hour then filtered using 0.22 μm Millex-GP PES membrane filters (Millipore SLGP033RS). The clarified lysate was loaded onto a HisTrap FF Crude column (Cytiva 29048631) pre-equilibrated with wash buffer (50 mM HEPES [pH 6.5], 400 mM NaCl, 1 mM MgCl_2_, 10% w/v glycerol, 20 mM imidazole, 5 mM 2-mercaptoethanol). The column was washed with 25 mL wash buffer before elution with elution buffer (wash buffer with 400 mM imidazole). The eluent was treated with TEV protease (~0.1 mg/ml) and dialyzed against 1 L of dialysis buffer (20 mM HEPES [pH 6.5], 100 mM NaCl, 1 mM MgCl_2_, 10% w/v glycerol, 5 mM 2-mercaptoethanol) for 3 hours at 4°C. The sample was then loaded onto a HiTrap SP HP column (Cytiva 29051324) pre-equilibrated with dialysis buffer, and eluted over a gradient of dialysis and elution (20 mM HEPES [pH 6.5], 10 M NaCl, 1 mM MgCl_2_, 10% w/v glycerol, 5 mM 2-mercaptoethanol) buffers. Relevant fractions were pooled, concentrated using a 50 kDa Amicon Ultra-4 centrifugal filter (Millipore UFC805008), filtered using a 0.22 μm Millex-GP PES membrane (Millipore SLGP033RS), and loaded onto the Superdex 200 Increase 10/300 GL column (Cytiva 28990944) in binding assay buffer (50 mM Tris [pH 7.5], 80 mM NaCl, 5 mM MgCl_2_, 5% w/v glycerol, 0.002% Tween-20, 1 mM DTT). Concentration was determined using the colorimetric Bradford assay (Bio-Rad 5000006).

Ytm1-GFP purification was carried out at 4° C. Cells were lysed using a Dounce homogenizer (Thomas Scientific) in ~25 mL of lysis buffer (50 mM HEPES [pH 7.5], 400 mM NaCl, 1 mM MgCl_2_, 10 % w/v glycerol, 20 mM imidazole, 5 mM 2-mercaptoethanol, 0.2 mM ATP, 1 mM PMSF, 3 U/mL benzonase, 1X Roche complete protease inhibitor without EDTA) per L of initial cell culture. The crude lysate was centrifuged at 55,000 rpm in a Type 70 Ti rotor (Beckman Coulter) for 1 hour then filtered using 0.22 μm Millex-GP PES membrane filters (Millipore SLGP033RS). The clarified lysate was loaded onto a HisTrap FF Crude column (Cytiva 29048631) pre-equilibrated with wash buffer (50 mM HEPES [pH 7.5], 400 mM NaCl, 1 mM MgCl_2_, 10% w/v glycerol, 20 mM imidazole, 5 mM 2-mercaptoethanol). The column was washed with 25 mL wash buffer before elution with elution buffer (wash buffer with 400 mM imidazole). The eluent was treated with TEV protease (~0.1 mg/ml) and dialyzed against 1 L of dialysis buffer (20 mM HEPES [pH 7.5], 100 mM NaCl, 1 mM MgCl_2_, 10% w/v glycerol, 5 mM 2-mercaptoethanol) for 3 hours at 4°C. The sample was then loaded onto a HiTrap Q HP column (Cytiva 29051325) pre-equilibrated with dialysis buffer, and eluted over a gradient of dialysis and elution (20 mM HEPES [pH 7.5], 10 M NaCl, 1 mM MgCl_2_, 10% w/v glycerol, 5 mM 2-mercaptoethanol) buffers. Relevant fractions were pooled, concentrated using a 50 kDa Amicon Ultra-4 centrifugal filter (Millipore UFC805008), filtered using a 0.22 μm Millex-GP PES membrane (Millipore SLGP033RS), and loaded onto the Superdex 200 Increase 10/300 GL column (Cytiva 28990944) in binding assay buffer (50 mM Tris [pH 7.5], 80 mM NaCl, 5 mM MgCl_2_, 5% w/v glycerol, 0.002% Tween-20, 1 mM DTT). Concentration was determined using the colorimetric Bradford assay (Bio-Rad 5000006).

### Mass photometry

All mass photometry data were taken using a Refeyn OneMP mass photometer (Refeyn Ltd). Movies were acquired for 6,000 frames (60 s) using AcquireMP software (version 2.4.0) and analyzed using DiscoverMP software (version 2.4.0, Refeyn Ltd), all with default settings. Proteins were measured by adding 2uL of stock solution (100 nM) to an 8 uL droplet of filtered phosphate buffered saline (Gibco 14190144). Contrast measurements were converted to molecular weights using a standard curve generated with bovine serum albumin (Thermo 23210) and Urease (Sigma U7752). The data were fitted by a Gaussian in MATLAB (Mathworks, Natick, Ma) to determine the measured molecular weight.

### Native mass spectrometry (nMS)

The purified protein samples were buffer-exchanged into nMS solution (150 mM ammonium acetate, 0.01% Tween-20, pH 7.5) using Zeba desalting microspin columns with a 40-kDa molecular weight cut-off (Thermo Scientific). The buffer-exchanged sample was diluted to 2 μM and was loaded into a gold-coated quartz capillary tip that was prepared in-house. The sample was then electrosprayed into an Exactive Plus EMR instrument (Thermo Fisher Scientific) using a modified static nanospray source (Olinares and Chait, 2020). The MS parameters used included: spray voltage, 1.22 kV; capillary temperature, 200 °C; S-lens RF level, 200; resolving power, 17,500 at m/z of 200; AGC target, 1 × 10^6^; number of microscans, 5; maximum injection time, 200 ms; in-source dissociation (ISD), 100 V; injection flatapole, 8 V; interflatapole, 7 V; bent flatapole, 5 −6 V; high energy collision dissociation (HCD), 10 V; ultrahigh vacuum pressure, 6 × 10^−10^ mbar; total number of scans, 100. Mass calibration in positive EMR mode was performed using cesium iodide. Raw nMS spectra were visualized using Thermo Xcalibur Qual Browser (version 4.2.47). Data processing and spectra deconvolution were performed using UniDec version 4.2 (Marty et al., 2015; Reid et al., 2019). The UniDec parameters used were m/z range: 2,000 – 7,000; mass range: 10,000 – 200,000 Da; sample mass every 1 Da; smooth charge state distribution, on; peak shape function, Gaussian; and Beta softmax function setting, 20.

### Size exclusion chromatography coelution assays

All coelution experiments were run in binding assay buffer (50 mM Tris [pH 7.5], 80 mM NaCl, 5 mM MgC_l2_, 5% w/v glycerol, 0.002% Tween-20, 1 mM DTT) using a Superdex 200 Increase 10/300 GL column (Cytiva 28990944). Conditions run were MIDAS-WT alone, MIDAS-Y4666R alone, Rsa4-GFP alone, Rsa4-GFP with MIDAS-WT, Rsa4-GFP with MIDAS-Y4666R, Ytm1-GFP alone, Ytm1-GFP with MIDAS-WT, and Ytm1-GFP with MIDAS-Y4666R. In all cases, the MIDAS protein was at 60 μM and the Rsa4-GFP or Ytm1-GFP was at 40 μM in 500 μL total. Proteins were mixed and dialyzed against binding assay buffer for three hours prior to injection onto the column. The eluent was collected in 250 μL fractions, which were analyzed both by SDS-PAGE and by GFP fluorescence. SDS-PAGE was performed using precast Novex 4-20% Tris-Glycine gels (Thermo Fisher XP04205BOX). Coomassie-stained gels were imaged using a LiCOR Odyssey system. Fluorescence measurements were made using a Synergy Neo Microplate reader (488 nm excitation, 528 nm emission).

### Microscale thermophoresis (MST)

Binding affinities between Mdn1 MIDAS-WT or MIDAS-Y4666R and Rsa4-GFP or Ytm1-GFP were measured using a Monolith NT.115 instrument (NanoTemper Technologies, Munich, Germany). All experiments were run in binding assay buffer (50 mM Tris [pH 7.5], 80 mM NaCl, 5 mM MgCl_2_, 5% w/v glycerol, 0.002% Tween-20, 1 mM DTT), matching the buffer that all proteins were stored in. Rsa4-GFP or Ytm1-GFP were diluted to 100 nM and filtered using a 0.22 μm centrifugal filter (Millipore UFC30GV00), and mixed 1:1 with MIDAS protein (initial stock 80 μM, prepared in 1:2 serial dilutions) leading to a final concentration of 50 nM. Proteins were mixed for 5-10 minutes at room temperature in the dark before being loaded into Monolith NT. 115 series premium capillaries (NanoTemper MO-K025). Measurements were performed using 40-60% excitation power and the medium MST intensity option within the default settings in MO.Control software (NanoTemper Technologies, Munich, Germany). The raw fluorescence data were exported to MO.Affinity software and converted to |ΔF_Norm_| by comparing the fluorescence before heating to the fluorescence 1.5 seconds after heating. |ΔF_Norm_| as a function of MIDAS concentration could only be fitted successfully for MIDAS-WT with Rsa4-GFP, and were fitted with a binding isotherm in MATLAB (Mathworks, Natick, Ma):

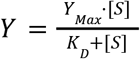

Where Y_max_ is the maximum system response. All MST measurements were repeated four times on four different days using at least two separate preparations of each of the proteins used.

### DNA substrates for optical tweezers force jump assay

The force jump assay tether consists of two 1.5 kilobase-pair biotinylated dsDNA handles connected by a ssDNA bridge. SNAP-GFP nanobody and SNAP-MIDAS were covalently attached to oligonucleotides that anneal to the bridge. The dsDNA handles are the same as were used in our previous study (Mickolajczyk et al., 2021). They were generated by PCR using primers designed to produce overhangs as needed to anneal the bridge stands. The forward primer for the 5’ overhang handle contains an inverted base (iInvd) that terminates Phusion DNA polymerase (with a sequence of 5’-CAACCATGAGCACGAATCCTAAACCT/ilnvdT/GCATAACCCCTTGGGGCCTCTAAACG-3’). The forward primer for the 3’ overhang handle contains inverted bases at its 5’ end that terminate Taq polymerase (with a sequence of 5’-/5InvdG//iInvdC//iInvdA//iInvdA//iInvdA//iInvdT//iInvdC//iInvdT//iInvdC//iInvdC//iInvdG//iInvdG//iInvdG//iInvdG//iInvdT//iInvdT//iInvdC//iInvdC//iInvdC//iInvdC//iInvdA//iInvdA//iInvdT//iInvdA//iInvdC//iInvdG/TAGTCTAGAGAATTCATTGCGTTCTGTACA/3ddC/-3’). The reverse primers for the 5’ and 3’ overhang handles contain a biotin at their respective 5’ ends for bead attachment. The ssDNA bridges, which anneal to the overhangs, were synthesized by Integrated DNA Technologies (Coralville, IA), The 55 nt bridge has the sequence 5’-GGGAGACAACCATGAGCACGAATCCTAAACCTCCTCACTGTCTCGTCCGTCGTTCCGTCCTGT<u>CCTTTCCCCTCTCTTTTTCTTTCCCCTTTCTCCTCTCTCCTCTCCCTTCTCTCCC</u>CTCACCTCACGTCCGCCAGATCCACAGTTCGTGTACAGAACGCAATGAATTCTCTAGACTA-3’, and the 70 nt bridge has the sequence 5’-GGGAGACAACCATGAGCACGAATCCTAAACCTCCTCACTGTCTCGTCCGTCGTTCCGTCCTGT<u>CCTTTCCCCTCTCTTTTTCTTTCCCCTTTCTCCTCTCTCCTCTCCCTTCTCTCCCCCCTTTCTCCTCTCT</u>CTCACCTCACGTCCGCCAGATCCACAGTTCGTGTACAGAACGCAATGAATTCTCTAGACTA-3’. In both cases the underlined region is the portion that remains single-stranded in the force jump assay.

The oligonucleotides used to connect SNAP tagged proteins to the bridge were also synthesized by Integrated DNA Technologies (Coralville, IA) and have the sequences: 5’-Am-<u>CTCCTCTCTTTT</u>ACAGGACGGAAGGACAGACGAGAAAGTGAG-3’ and 5’-GAACTGTGGATCTGGCGGACGTGAGGTGAG<u>TTTCTCCTTTCT</u>-Am-3’, where the underline regions are the portions that do not anneal to the bridge, and Am denotes modification with a free amino group. A benzylguanine (BG) group was covalently attached to the amino-oligonucleotides using BG-GLA-NHS (New England BioLabs S9151S) under manufacturer-suggested reaction conditions, and subsequently purified using Micro Bio-Spin P-6 gel columns (Bio-Rad 7326221). The BG-oligos were then conjugated to SNAP-proteins by mixing at a 2:1 molar ratio in storage buffer (20 mM HEPES [pH 7.5], 150 mM NaCl, 1 mM MgCl_2_, 3% w/v glycerol, 1 mM DTT) incubating in the dark at 4° C overnight. Excess oligonucleotides and unreacted SNAP proteins were removed by purification on a Superdex 200 Increase 10/300 GL column (Cytiva 28990944) in storage buffer. Fractions with oligo-protein conjugates were identified via SDS-PAGE, pooled, and flash frozen in liquid nitrogen prior to use.

To prepare for experiments, the 5’ overhang dsDNA handle and bridge strands were first annealed by mixing 50 nM of 5’ overhang handles with 12.5 nM of bridge ssDNA in storage buffer, incubating at 65° C for 1 minute, then cooling to 4° C at 0.1° C/s. This annealed product was then mixed with oligo-SNAP-nanobody and oligo-SNAP-MIDAS (final 2 nM, 10 nM, 10 nM, respectively) in optical tweezers assay buffer (50 mM Tris [pH 7.5], 80 mM NaCl, 5 mM MgCl_2_, 0.1 mg/mL BSA, 0.002% Tween-20, 1 mM DTT) and incubated on ice for 30 minutes. This reaction was then diluted 5-fold into optical tweezers assay buffer containing 0.08% w/v streptavidin-coated polystyrene beads (2.1 μm diameter, Spherotech) and incubated on ice for at least 15 minutes. This mixture was diluted 1,000-fold in optical tweezers assay buffer at the time of use. Separately, 5 nM 3’ overhand handles were mixed with 0.08% w/v streptavidin-coated polystyrene beads (2.1 μm diameter, Spherotech) in optical tweezers assay buffer, and incubated on ice for at least 15 minutes. This mixture was also diluted 1,000-fold in optical tweezers assay buffer at the time of use.

To form a tether, a bead conjugated to the 5’ overhang handle complex was immobilized on a pipette via suction and brought into close proximity to a second bead conjugated to 3’ overhang handles held in an optical trap. Upon hybridization of the 3’ overhang handle with the 5’ overhang handle complex, a tether was formed. For both the 55 and 70 nt bridge constructs, we ensured that only a single “tether” was drawn between each pair of beads by measuring the force-extension curve and matching it to a well-characterized hairpin construct held between identical dsDNA handles (**Figure 2D**) (Mickolajczyk et al., 2021).

### Optical tweezers force jump assay

The force jump assay developed for this study is related to the ReaLiSM and junctured-DNA-tweezers approaches (Kim et al., 2010; Kostrz et al., 2019). All single-molecule measurements were made using an optical tweezers instrument (“MiniTweezers”) (Smith et al., 2003). Optical tweezers assay buffer (50 mM Tris [pH 7.5], 80 mM NaCl, 5 mM MgCl_2_, 0.1 mg/mL BSA, 0.002% Tween-20, 1 mM DTT) was used for all experiments. At the start of each experiment, a tether was formed in the sample chamber and its force-extension behavior was obtained by moving the optically trapped bead at a speed of 70 nm/s away from the second bead held in the pipette. Data collected from the tether was only accepted if the force-extension curve could be overlaid on that of a previously characterized tether of comparable length containing a hairpin (Mickolajczyk et al., 2021). Rsa4-GFP or Ytm1-GFP were injected using a shunt line to a final of 20 nM before each tether was pulled. The experiment was run by using constant force mode and switching between 0.5 pN low force and 4-12 pN high force. All force levels were held for at least five seconds (empirically determined to be long enough to include all events). Force feedback was temporarily turned off at the low force to prevent large position fluctuations. Data was collected at 200 Hz (instrument response time ~5-10 ms). All experiments were run at 23 ± 1 °C.

Single-molecule data were analyzed in MATLAB (Natick, MA). High-force positional plateaus were investigated using a two-state hidden Markov model, which was modified from existing code (Mickolajczyk et al., 2015). The data were centered at 0 by subtracting the mean of the first five data points from the whole vector, and the effective standard deviation was determined by building a distribution of 10-data point boxcar standard deviations (eSD), and finding the median. The intermediate (proteins bound) and final (proteins unbound) positions on the high-force plateau were treated as Gaussian emitters, with centers zero and + 4 eSD units, and standard deviations one eSD unit. A transition matrix was constructed to only allow transitions from the zero-position state to the higher position state. The emission matric, transition matrix, and experimental data were used as input to the Viterbi algorithm (Viterbi, 1967), which returned the most likely sequence of hidden states. If a state change was detected, then the bond lifetime was calculated as the duration preceding the state change, and Δx was calculated as the difference in means between an equal number of data points (corresponding to the binding duration) before and after the state change. In some cases, there was drift in the data, which could lead to a larger position change than the Δx jump over time. In these cases, the high-force plateau was truncated to the area just around the Δx jump. Events were only accepted if they were at least four data points long. Every event was manually inspected, and only events that were clearly an instantaneous change in motion (rather than drift) were retained for further analysis. Pooled values of bond lifetime data were fitted to a single-exponential:

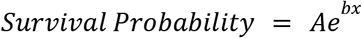

### Mechanical circuit model

All spring elements in the mechanical circuit model were modeled using the worm-like chain model:

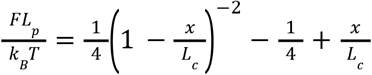

Where L_c_ is the contour length and L_p_ is the persistence length. The parameters used were persistence length 1 nm, and contour length 0.59 nm per DNA nucleotide (Liphardt et al., 2001). The bridge strand was 55 or 70 nt in length, and the loading strand was 24 nt in length. When both springs were engaged (proteins bound), the force-extension curve of the system was:

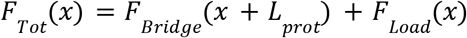

When the proteins dissociated, the force-extension curve of the system was:

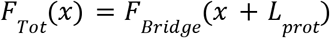

Where L_prot_ is the inextensible length of the proteins. Based on existing structural data, we estimate L_prot_ to be 8 nm (Ahmed et al., 2019). For a given total force placed on the system (F_Tot_), the force placed on the proteins (F_Load_) and the theoretical Δx could be calculated. All data analysis and fitting was performed in MATLAB (Natick, Ma).

## SUPPLEMENTAL INFORMATION

Four supplemental figures

## ACKNOWLEDGEMENTS

We thank members of the Kapoor and Liu Labs for useful discussions. We thank Jun Funabiki and the Paul Nurse Lab for generously providing the *S. pombe* mitotic cDNA and Xiaocong Cao (Liu Lab) for providing DNA handle reagents and protocols. We thank the Rockefeller High-Throughput Screening Resource Center for assistance with the MST experiments. T.M.K is grateful to the NIH/NIGMS for funding (GM98579 and R35 GM130234-01). B.T.C. is grateful to NIH/NIGMS for funding (P41 GM109824 and P41 GM103314). S.L. is supported by the Robertson Foundation and NIH (DP2HG010510). K.J.M. is supported by a National Cancer Institute K00 Fellowship (K00CA223018).

## AUTHOR CONTRIBUTIONS

K.J.M. and T.M.K conceived of the project. K.J.M. designed and carried out experiments, made plasmids, purified and labeled proteins, and analyzed data. P.D.O. performed native mass spectrometry experiments with supervision by B.T.C. S.L. provided optical tweezers access, materials, and expertise. K.J.M. and T.M.K. wrote the manuscript with contributions from all other authors.

