## Supplementary material for "The MIDAS domain of AAA mechanoenzyme Mdn1 forms catch bonds with two different substrates": Supp Material

**This PDF file includes:**

**Figs. S1 to S4**

### Supplementary Figures

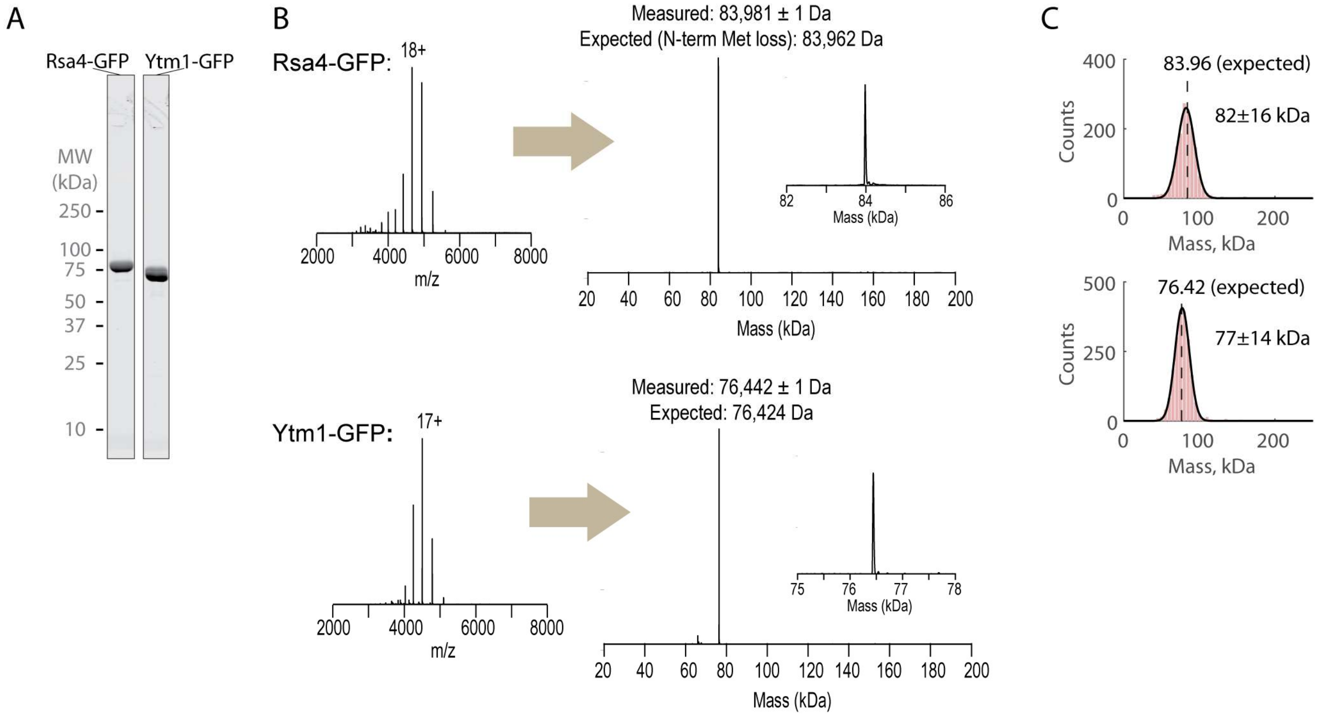


**Figure S1. Preparations of recombinant Rsa4-GFP and Ytm1-GFP**
**(A)** SDS-PAGE gels (Coomassie staining) of final purified Rsa4-GFP and Ytm1-GFP.
**(B)** Raw and deconvolved native mass spectrometry spectra for Rsa4-GFP and Ytm1-GFP.
**(C)** Mass photometry measurements of Rsa4-GFP (top, 1,510 measured molecules) and Ytm1 (bottom, 2,031 measured molecules). Measured mass shown inset at mean ± standard deviation (SD) of fitted Gaussian (black line).

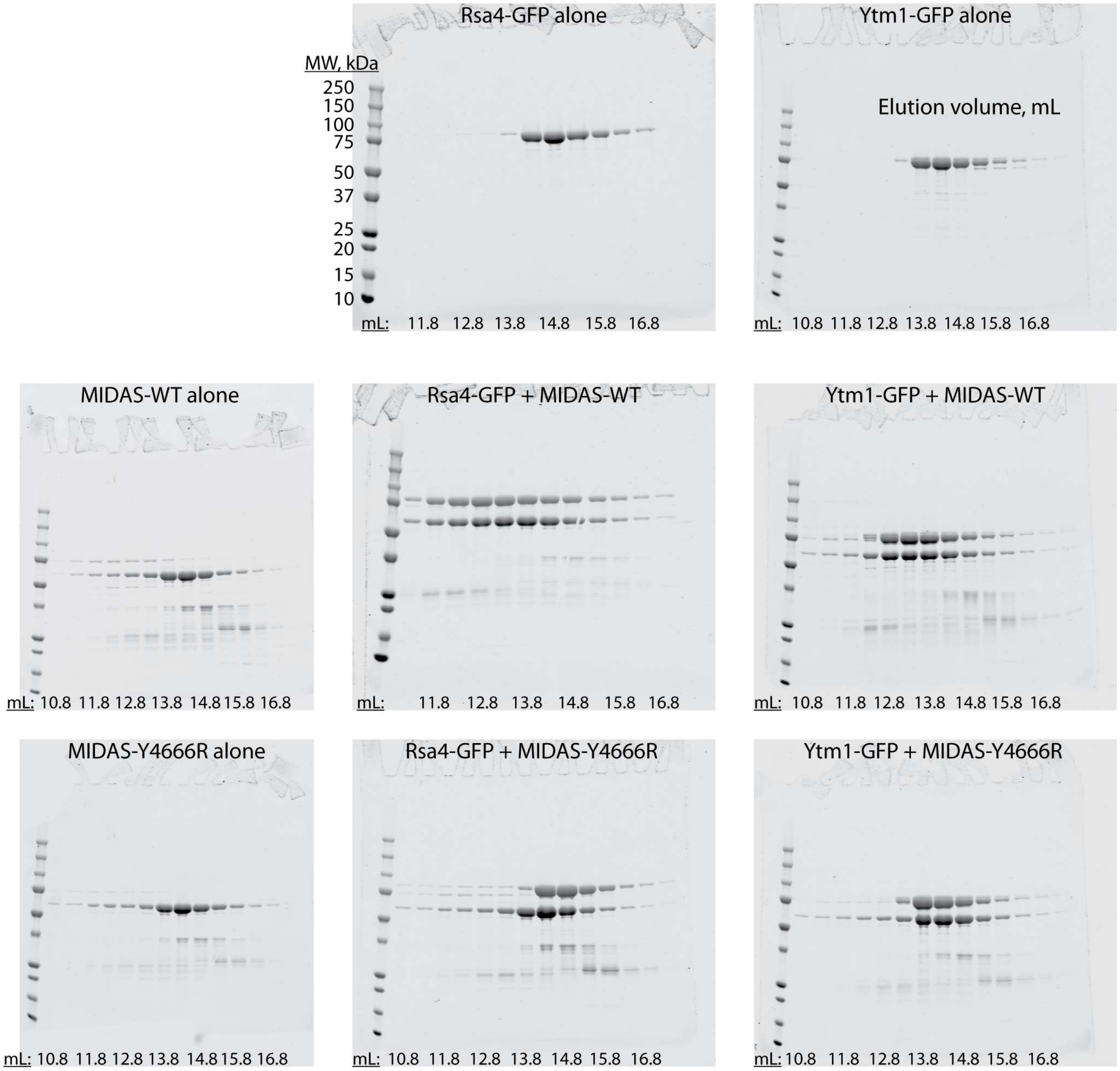


**Figure S2. Full gels for size exclusion chromatography experiments.**
Uncropped SDS-PAGE gels (Coomassie staining) showing size exclusion chromatography results for the listed proteins. The same molecular weight ladder was used for all gels. Corresponds to **Figure 1 E-F** and **Figure 4 B-C** in the main text.

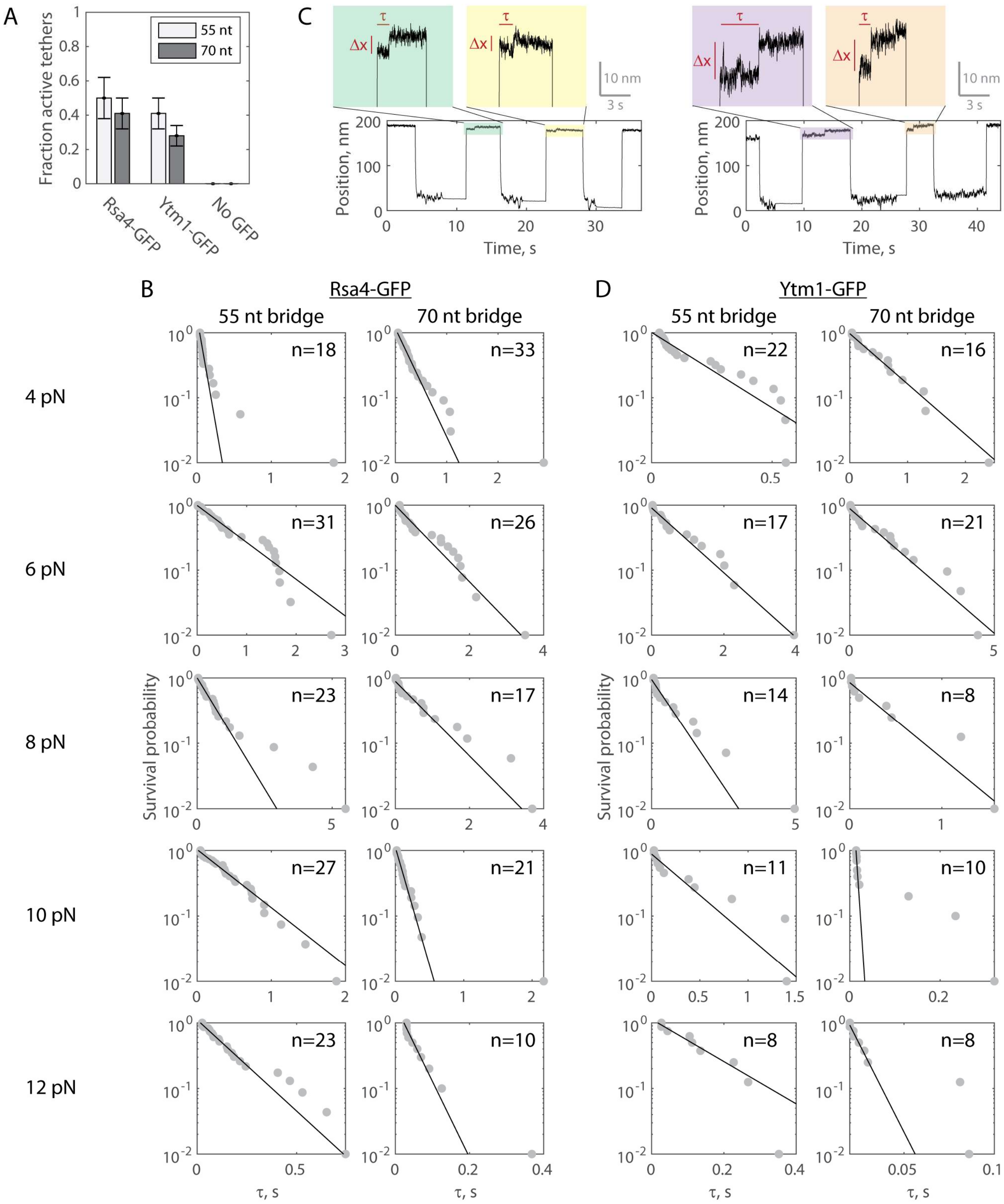


**Figure S3. Full datasets generated in the force jump assay**

**(A)** Fraction of verified single tethers that displayed Δx events with Rsa4 present (26/52 and 27/66 on the 55 and 70 nt bridge constructs, respectively), Ytm1 present (33/80 and 24/86), or no GFP protein present (0/13 and 0/22). At least 20 jumps and two different high force levels were tested for each tether. Data pooled from at least three independent experimental days and shown as total with propagated Poisson counting error.

**(B)** Full dataset (at each $F_{Tot}$) for force jump assay on both bridge constructs with Rsa4-GFP. Data in gray, fitted single exponential in black. The number of measurements (n) labeled inset for each experiment. For all distributions, the final data point was moved from y=0 to y=0.01 to enable semilog plotting. Black lines show fits to a single exponential.

**(C)** Example force jump data on the 55 nt bridge (left) and 70 nt bridge (right) with Ytm1-GFP (20 nM) added.

**(D)** Full dataset for force jump assay on both bridge constructs with Ytm1-GFP. Data in gray, fitted single exponential in black. The number of measurements (n) labeled inset for each experiment.

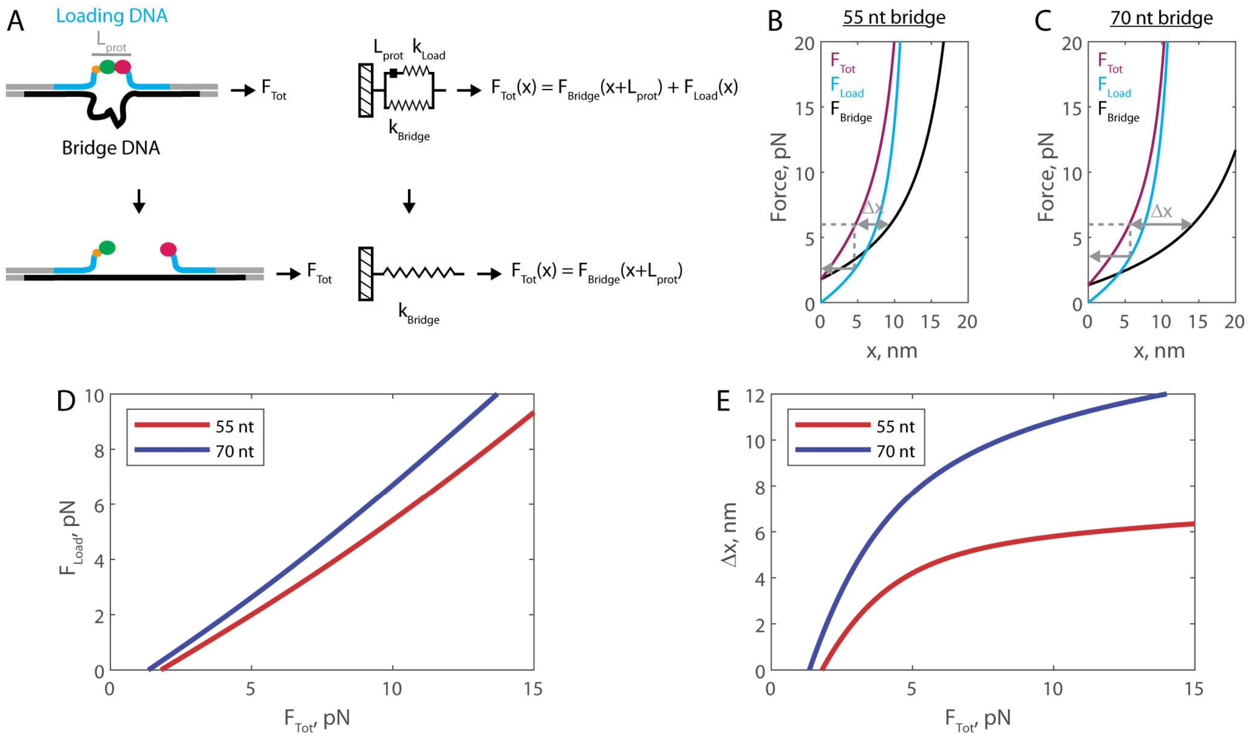


**Figure S4. Mechanical circuit model for the force jump assay**

**(A)** Mechanical circuit model describing the force jump assay. When the proteins (of inextensible length $L_{prot}$) are bound, force is partitioned between the top "loading" strand (two 12 nt single-stranded regions) and the bottom bridge strand. In the equations shown inset, x designates extension along the loading stand spring.

**(B)** Modeled force-extension curves for the force jump assay using the 55 nt bridge construct. Gray lines show an example for 6 pN total applied force: the single-headed arrow shows the amount of force felt by the proteins, and the double-sided arrow shows the expected magnitude of Δx when the proteins dissociate.

**(C)** Similar to panel B, using the 70 nt bridge construct.

**(D)** Transfer function showing the calculated $F_{Load}$ for a given $F_{Tot}$.

**(E)** Transfer function showing the calculated Δx for a given $F_{Tot}$.
